# Mind the gap: Understanding discordance between culture- and a non-culture-based measure of bacterial burden in murine tuberculosis treatment models

**DOI:** 10.64898/2025.12.18.695164

**Authors:** Samuel T. Tabor, Allan D. Friesen, Matthew J. Reichlen, Christian Dide-Agossou, Max McGrath, Ryan Peterson, Vitaly V. Ganusov, Gregory T. Robertson, Martin I. Voskuil, Nicholas D. Walter

## Abstract

The standard pharmacodynamic marker in murine tuberculosis drug studies is colony-forming units (**CFU**). A faster PCR-based marker of bacterial burden is 16S rRNA. For unclear reasons, treatment reduces CFU more than 16S rRNA. We evaluated this CFU-16S gap and estimated the fraction potentially attributable to slow decay of 16S rRNA from dead *Mycobacterium tuberculosis* (***Mtb***) versus transition to a viable but not culturable on solid agar (**VBNC_SA_**) population.

We quantified the CFU-16S gap during and following treatment in six BALB/c mouse studies and one *in vitro* study. Applying a two-population ordinary differential equation-based model of *Mtb* death and 16S rRNA decay, we estimated the fraction of the gap potentially attributable to dead *Mtb.* Using meta-regression, we estimated the association between CFU or 16S rRNA with relapse.

For all regimens, CFU fell more than 16S rRNA, ranging from isoniazid-rifampin-pyrazinamide-ethambutol (CFU decreased 39-times more than 16S rRNA at week 4) to bedaquiline-pretomanid-moxifloxacin-pyrazinamide (CFU decreased >500,000-times more). The two-population model suggested that the fraction of the CFU-16S gap attributable to residual 16S rRNA from dead *Mtb* is modest and decreases over time. After treatment, 16S rRNA often fell while CFU rose. Four-week CFU change explained most variation in relapse (R²=0.90) while four-week 16S rRNA change did not (R²=0.24).

CFU-16S gap is only partially explained by slow decay of residual 16S, suggesting development of a VBNC_SA_ population. However, continued decrease in 16S rRNA after treatment cessation and its limited association with relapse suggests VBNC_SA_ may be a transient rather than persistent state.

## INTRODUCTION

Tuberculosis (**TB**) is the leading cause of death from a single infectious agent and among the top 10 causes of death worldwide.^1^ There is an urgent need for shorter, more efficacious TB treatments. Murine studies are a cornerstone of preclinical drug and regimen evaluation that strongly influence which regimens are advanced to human testing.^2–4^ In murine models, drug effects are assessed based on change in pharmacodynamic (**PD**) markers during treatment and/or on the occurrence of relapse after treatment completion.^5–8^

For decades, the traditional readout of *M. tuberculosis* (***Mtb***) burden in murine TB treatment models has been colony-forming units (**CFU**) that grow when mouse lung homogenate is cultured on solid agar.^5–8^ More recently, a proposed non-culture readout of bacterial burden is *Mtb* 16S rRNA measured by qPCR. *Mtb* 16S rRNA has been evaluated as an alternative PD marker in both human sputum^9–17^ and murine lung homogenate.^18–20^ In this work, we address a gap in understanding the relationship between the two PD markers of bacterial burden in the conventional BALB/c high-dose aerosol infection model that is a reference standard in preclinical drug evaluation. Our focus here is exclusively on 16S rRNA in preclinical rather than human studies.

*Mtb* 16S rRNA was originally proposed as a measure of bacterial burden that could accelerate drug and regimen evaluation by providing similar information as CFU, more rapidly.^9,18,11^ However, recent murine studies^18,19^ have identified a currently unexplained “gap” between CFU and 16S rRNA measurements. Specifically, CFU and 16S rRNA numbers dissociate during treatment with certain individual drugs and regimens with CFU typically declining faster than 16S rRNA levels.^18,19^ Two non-exclusive explanations have been proposed for why CFU may decrease more than 16S rRNA. First is the development of an *Mtb* population that is viable but non-culturable on solid agar (**VBNC_SA_**). It is well established that culture on solid agar (*i.e.,* CFU) may not recover all viable *Mtb.*^18,19^ By contrast, qPCR should amplify 16S rRNA irrespective of whether *Mtb* is capable of growth on agar, thereby measuring both culturable and non-culturable bacteria. If drug treatment causes a large share of the *Mtb* population to shift to a VBNC_SA_ state, 16S rRNA might fall to a lesser degree or more slowly than CFU. A second explanation is detection of residual 16S rRNA from dead *Mtb.*^6,21^ It is unclear whether and for how long 16S rRNA remains quantifiable in dead *Mtb.* If 16S rRNA remains persistently detectable after bacterial death, 16S rRNA might fall to a lesser degree or more slowly than CFU.

Here, we aimed to clarify interpretation of PD results in murine TB drug evaluation by addressing several questions. First, how commonly and under what circumstances is the gap between CFU and 16S rRNA observed in murine studies? We addressed this by evaluating CFU and 16S rRNA in (1) untreated mice following low-dose aerosol infection, (2) mice treated with diverse drugs in monotherapy, (3) mice treated with diverse regimens, and (4) mice allowed a drug -free holiday to recover after non-curative drug treatment. Second, we conducted an *in vitro* study that replicated the gap between CFU and 16S rRNA. Finally, leveraging three recent relapsing mouse model (**RMM**) studies, we conducted modeling to determine if CFU count or *Mtb* 16S rRNA better predicted treatment time required to prevent relapse in 95% (T_95_) of mice^6^ and asked if a combination of the two PD markers might better explain variation in T_95_.

## METHODS

### Murine studies

Supplemental Information includes full details of each experiment. Common elements are briefly described here. Female BALB/c mice 6-8 weeks old were infected via high-dose (∼10^4^ CFU/mouse) or low-dose (∼10^2^ CFU/mouse) aerosol with *Mtb* Erdman at Colorado State University (CSU) or *Mtb* H37Rv at Johns Hopkins University (JHU). Mice were treated with individual drugs or regimens via oral gavage and humanely euthanized at time points described in Supplemental Information. Lung lobes were aseptically dissected and flash frozen in liquid nitrogen prior to determination of CFU and 16S rRNA. All animal procedures were conducted according to relevant national and international guidelines and approved by institutional animal safety committees as described in the Supplemental Information.

### In vitro HRZE and H_2_O_2_ exposure

*Mtb* Erdman was cultured *in vitro* using Middlebrook 7H9 broth (Difco) supplemented with 0.085 g/l NaCl, 0.2% glucose, 0.2% glycerol, 0.5% BSA, and 0.05% Tween-80 at 36.5°C and 5.0% CO_2_. Single use frozen *Mtb* aliquots were revived in 7H9 and grown to mid-log phase then cultures were diluted to OD_600_=0.05, dispensed in 5.0 ml aliquots into sterile glass tubes (20 by 125 mm) containing sterile stir bars (12 by 4.5 mm), and outgrown for 18 h under rapid agitation (∼200 rpm stirring speed using a rotary magnetic tumble stirrer) prior to the initiation of drug exposure. RNA was collected after exposure to isoniazid (H), rifampin (R), pyrazinamide (Z), ethambutol (**HRZE**) (0.5 µg/mL of H, 0.5 µg/mL of R, 20 µg/mL of E, and 16 µg/mL of Z) and 200 mM H₂O₂, respectively, as further described in Supplemental Information.

### Laboratory analysis

In murine studies, CFU was measured by serial dilution of lung homogenates and subsequent plating on 7H11-oleic acid-albumin-dextrose-catalase (OADC) agar supplemented with 0.4% activated charcoal. Following RNA extraction, reverse transcription and 1:10 dilution of cDNA, absolute 16S rRNA copies were quantified via TaqMan quantitative PCR (qPCR) as detailed in Supplemental Information.

### Statistical analysis

#### Inferential statistics

Pairwise relationships between measures were investigated using Pearson correlation coefficients and their associated significance tests. Wilcoxon rank-sum tests were used to determine differences between single drugs and between regimens, respectively.

#### Definition of the gap

During treatment, *Mtb* CFU declined more rapidly than 16S rRNA counts, resulting in an apparent excess of 16S rRNA relative to CFU. We quantified the *logarithmic gap* after treatment time *t* as the difference in logarithms (base 10) of change between CFU and 16S rRNA counts relative to pre-treatment values (**Fig 2a**). Specifically, the gap at time *t* after treatment is:

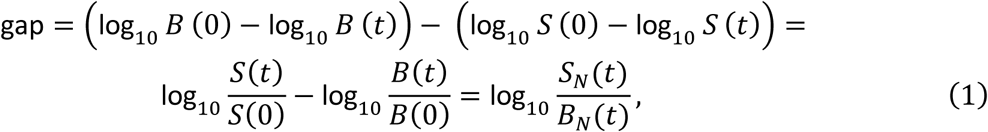

where *B*(0) and *S*(0) are CFU and 16S rRNA at the start of treatment, respectively, and *B*_*N*_(*t*) = *B*(*t*)/*B*(0) and *S*_*N*_(*t*) = *S*(*t*)/*S*(0) are the CFU and 16S rRNA at time *t* normalized to their values at the start of treatment. Note, if CFU were undetectable (i.e., *B*(*t*) = 0), the gap is undefinable.

Therefore, for the few observations where CFU=0, we set the CFU value *B* = 0.1. A gap of zero indicates that the proportionate decline of CFU and 16S rRNA are equal. A positive gap indicates that 16S rRNA declined more slowly than CFU, producing an apparent excess of rRNA relative to CFU. A negative gap indicates that 16S rRNA declined more rapidly than CFU.

### Attribution of gap to residual 16S rRNA from dead Mtb

Using mathematical modeling, we asked what percentage of the gap might be attributable to slow decay of 16S rRNA from already-killed *Mtb* while assuming a range of 16S rRNA decay rates from dead *Mtb*. Specifically, we propose a two-population ordinary differential equation (**ODE**)-based model of bacteria death and decay of 16S rRNA that we call the “Live-Dead” (**LD**) model (**Fig. 2b**). In this model, live/plateable bacteria (denoted *B*) die at per capita rate δ_*B*_ (half-life time = ln(2) /δ_B_) during treatment, becoming dead bacteria *D*. 16S rRNA from dead bacteria decay at rate δ_S_(half-life time = ln(2) /δ_S_):

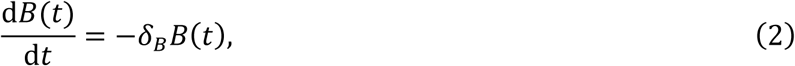

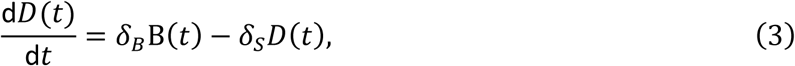

where the total 16S rRNA copies in the population *S*(*t*) are the sum of rRNA coming from live and dead bacteria:

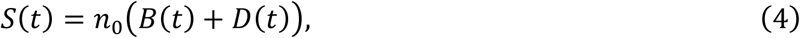

and *n*_0_is the number of 16S rRNA copies per cell. For simplification, our model assumes *n*_0_ is constant over time and is the same for live and dead bacteria, and there is no bacterial replication during treatment. Additionally, the model assumes that 16S rRNA comes only from live and dead bacteria (*i.e.*, the model does not allow for the possibility of VBNC_SA_ bacteria). Once a large population of dead bacteria accumulates during treatment, the total counts of 16S rRNA in this model decline exponentially:

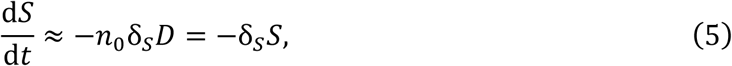

where during effective treatment most 16S rRNA during treatment comes from dead bacteria (*B* ≪ *D*). Thus, loss of 16S rRNA in the LD model is governed by the decay rate δ_*S*_. Note that to describe bi-exponential decline in CFU counts in our *in vitro* experiments (**Fig 2a**) we assumed that viable population *B* consists of two sub-populations *B*_1_and *B*_2_ (*B* = *B*_1_ + *B*_2_) with different decay rates δ_*B*1_ and δ_*B*2_. To reduce the number of fitted parameters, we fixed the initial values of viable bacteria (*B*(0)) and total 16S rRNA counts (*n*_0_*B*(0)) predicted by the model to the geometric mean values in the data.

To calculate the percent of the gap between observed CFU and 16S rRNA counts (*e.g.,* **Fig 2a**) that may be due to residual 16S rRNA from dead bacteria in the LD model we use the following formula:

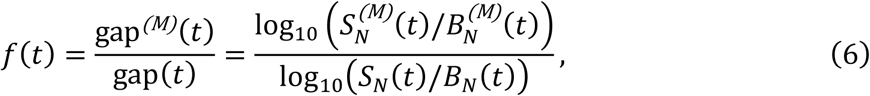

where 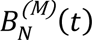 and 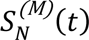 are the CFU and 16S rRNA numbers predicted by the LD model and normalized to their initial values (see **eqn. (1)** and **Fig 2c**). We calculated the *f(t)* as the proportion of the gap between CFU and 16S rRNA counts as explained by the LD model (**eqns. (2)-(4)**) assuming different decay rates (or half-life times) of the 16S rRNA in dead bacteria δ_*S*_ (see **eqn. (5)**). Note that the fraction of gap (**eqn (6)**) in principle can be larger than 100% if 16S rRNA decays at a slower rate than that observed experimentally.

### Half-life estimation from in vitro H_2_O_2_ exposure

To determine a range of plausible half-life values for 16S rRNA from dead *Mtb in vitro*, we evaluated decrease in 16S rRNA counts over time after rapid killing with high-concentration H_2_O_2_. Three biological replicates were collected per time point. For each time point, the median 16S rRNA across the three replicates was used as the central estimate. Analysis was limited to days 0–7, with day 7 being the last time point at which 16S rRNA remained detectable. Interval-specific decay rates between consecutive days were computed under a first-order decay assumption using the equation*: k = -Δ*ln*(C) / Δt* where *C* is the median 16S rRNA and *Δt* is the elapsed time between measurements. The half-life for each interval was then calculated as: T_1/2_ = ln*(2) / k* and converted to hours. An overall decay rate over the 0–7 day window was estimated by ordinary least squares regression of ln(median 16S rRNA) versus time; the corresponding half-life was computed as: T_1/2_ = ln(2) / (-slope).

#### Modeling of relapse (T_95_)

To determine whether CFU, 16S rRNA or a combination of both better explains variation in relapse, we used results of three already-completed BALB/c RMM studies that tested 9 unique regimens as described in Supplemental Information. Using established methods,^7^ we first estimated T_95_ (*i.e.,* the treatment duration projected to cure 95% of mice) for each drug regimen via a Bayesian sigmoidal Emax model using the function “stan_emax” in the rstanemax R package, as further described in Supplemental Information.^6,7,22^

#### Modeling contribution of changes in CFU and 16S rRNA to predict T_95_ estimates using meta-regression

To determine how changes in CFU and 16S rRNA from baseline to day 28 predict observed T_95_ estimates, we performed a meta-regression analysis weighted by the inverse standard error of T_95_ estimates, fitting three models using maximum likelihood estimation. The first model included baseline and day 28 changes in CFU only; the second model used baseline and day 28 changes in 16S rRNA only; and the third incorporated changes in both CFU and 16S rRNA. Models were compared against using pseudo-R-squared, Akaike Information Criterion (AIC), Root Mean Square Error (RMSE), and likelihood ratio tests (each model was tested against a null model and a full model). Analysis was conducted using R (v 4.5.1; R Development Core Team, Vienna, Austria).

## RESULTS

### Minimal gap between CFU and 16S rRNA in untreated mice

In the BALB/c “chronic” model in which mice are infected via low-dose aerosol and then followed over time without treatment (Experiment 1), log_10_ CFU and log_10_ 16S rRNA changed concordantly (R² = 0.92, correlation coefficient = 0.96, *P =* 3.4×10^-14^, slope = 0.83)) (**Fig 1a-b**), 92% of the variation in CFU was explained by 16S rRNA change, confirming previous observations that 16S rRNA is an excellent, albeit nonlinear, correlate for CFU for CFU in the absence of drug treatment.^18^ As expected in the chronic model,^23–25^ the burden of *Mtb* measured by CFU or 16S rRNA stabilized after the onset of adaptive immunity 14-20 days after infection.^26^

**Fig 1.**
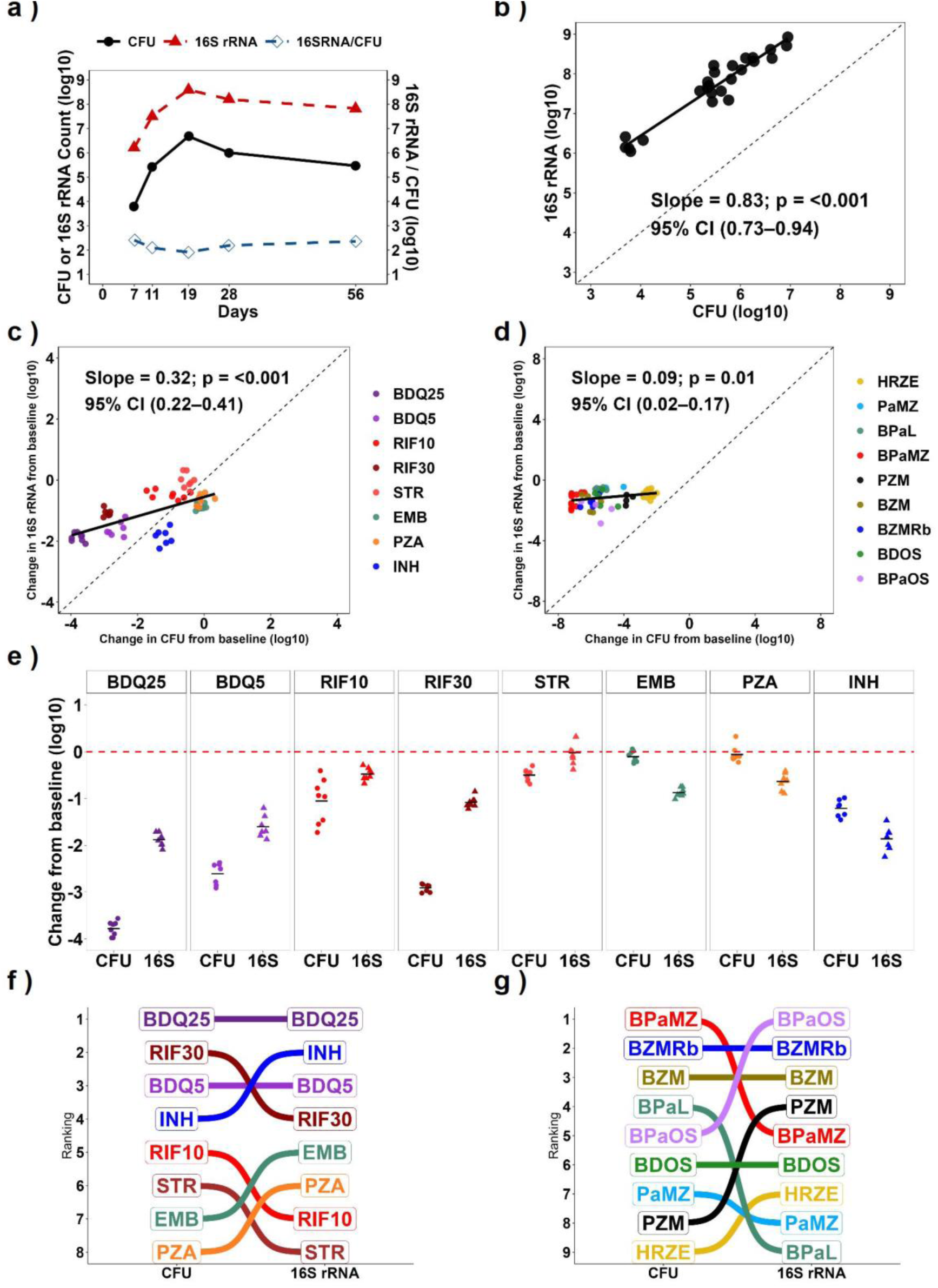
Relationship between 16S rRNA and CFU. **a.** CFU and Mtb 16S rRNA in the lungs of untreated BALB/c mice following low-dose aerosol infection in Experiment 1, with their ratio (16S rRNA/CFU). Circles indicate the mean CFU values, triangles indicate the mean 16S rRNA values, and lines connect mean values at each time point. **b.** Correlation between lung CFU and 16S rRNA in Experiment 1. **c.** Correlation between lung CFU and 16S rRNA in mice treated with individual drugs for four weeks in the high-dose aerosol infection model in Experiment 2. **d.** Correlation between lung CFU and 16S rRNA in mice treated with regimens for four weeks in Experiments 3–5. In panels b–d, circles indicate individual mice, and the line represents the fitted slope with its coefficient. **e.** Log10 reduction in lung CFU and 16S rRNA relative to pre-treatment controls in Experiment 2. Circles and triangles indicate CFU and 16S rRNA reductions for individual mice, respectively; horizontal bars indicate means, and the red dashed line marks the pre-treatment control reference. **f.** Ranking of individual drugs in Experiment 2 based on CFU and 16S rRNA. **g.** Ranking of regimens in Experiments 3–5 based on CFU and 16S rRNA.

### Variable gap between CFU and 16S rRNA in mice treated in monotherapy

In the BALB/c mouse high-dose aerosol infection model, 4-week treatment with six individual drugs decreased CFU and 16S rRNA to varying degrees (Experiment 2), resulting in lower correlation between CFU and 16S rRNA (0.64) than that observed for untreated mice (**Fig 1c**). The R² was 0.42, indicating only 42% of the variation in CFU was explained by 16S rRNA change. Additionally, the regression slope between CFU and 16S rRNA also decreased (slope = 0.34). The gap between CFU and rRNA was drug-specific. For certain drug exposures, CFU decreased more than 16S rRNA (**Table 1**). For example, bedaquiline (B) 25 mg/kg reduced CFU by 3.8 log_10_ but reduced 16S rRNA by 1.9 log_10_, producing a gap of 1.9 log that corresponds to an 80-times greater reduction in CFU than in 16S rRNA (**Fig 1e**). Rifampin 30mg/kg reduced CFU by 2.9 log_10_ but reduced 16S rRNA by 1.10 log_10_, producing a gap of 1.8 log that corresponds to a 67-times greater reduction in CFU than in 16S rRNA. Conversely, for other drug exposures, 16S rRNA decreased more than CFU. For example, isoniazid reduced CFU by 1.2 log_10_ but reduced 16S rRNA by 1.9 log_10_, producing a gap of −0.7 log that corresponds to a 4.5-times smaller reduction in CFU than in 16S rRNA. Ethambutol reduced CFU by only 0.1 log_10_ but reduced 16S rRNA by 0.9 log_10_, producing a gap of −0.8 log that corresponds to a 5.9-times smaller reduction in CFU than in 16S rRNA.

**Table 1.** Mean Log_10_ change in lung CFU and 16S rRNA relative to pre-treatment control after four weeks of treatment with individual drugs in the BALB/c high-dose aerosol model.

| Regimen | Change from PreRx to week 4 |  | Gap<sup>a</sup> | Reduction in CFU relative to reduction in 16S rRNA |
| --- | --- | --- | --- | --- |
|  | CFU (log<sub>10</sub>) | 16S rRNA (log<sub>10</sub>) |  |  |
| Bedaquiline 5mg/kg | -2.61 | -1.60 | 1.00 | 10.1-times greater reduction in CFU |
| Bedaquiline 25mg/kg | -3.78 | -1.88 | 1.90 | 79.8-times greater reduction in CFU |
| Rifampin 10mg/kg | -1.05 | -0.48 | 0.57 | 3.7-times greater reduction in CFU |
| Rifampin 30mg/kg | -2.91 | -1.08 | 1.83 | 67.0-times greater reduction in CFU |
| Streptomycin | -0.50 | -0.02 | 0.48 | 3.0-times greater reduction in CFU |
| Isoniazid | -1.21 | -1.86 | -0.65 | 4.5-times smaller reduction in CFU |
| Pyrazinamide | -0.06 | -0.64 | -0.57 | 3.8-times smaller reduction in CFU |
| Ethambutol | -0.10 | -0.88 | -0.77 | 5.9-times smaller reduction in CFU |

a. Negative Gap indicates more decline in 16S rRNA than CFU

CFU and 16S rRNA provided different readouts of the relative activity of drugs. For example, bedaquiline reduced CFU significantly more than isoniazid (*P*=0.001) but the effects of bedaquiline and isoniazid on 16S rRNA were indistinguishable (*P=*1). Streptomycin reduced CFU significantly more than ethambutol (*P*<0.001), but ethambutol reduced 16S rRNA more than streptomycin (*P*=0.0002). Pairwise comparisons between the other individual drugs for both CFU and 16S rRNA are shown in **Supplementary Table** S6. The rank order of drug effect differed if quantified based on CFU or 16S rRNA. Although both CFU and 16S rRNA ranked bedaquiline 25mg/kg as being the most efficacious, the ranking of other drugs changed. For instance, ethambutol tied with pyrazinamide for least effect on CFU but had the fifth greatest effect on 16S rRNA. Isoniazid had the fourth greatest effect on CFU but the second greatest effect on 16S rRNA (**Fig 1f**).

Both CFU and 16S rRNA identified dose-effect relationships for rifampin and bedaquiline but this relationship was more pronounced with CFU than 16S rRNA (**Table 1**). Relative to rifampin 10mg/kg, rifampin 30mg/kg resulted in a 1.8 log_10_ greater decrease in CFU but only a 0.6 log_10_ greater decrease in 16S rRNA. Similarly, relative to bedaquiline 5mg/kg, bedaquiline 25mg/kg resulted in a 1.2 log_10_ greater decrease in CFU but only a 0.3 log greater decrease in 16S rRNA.

### Large gap between CFU and 16S rRNA in mice treated with drug regimens

Evaluation of mice treated with 9 unique regimens in three RMM studies (Experiments 3-5) identified a lower correlation between CFU and 16S rRNA than observed in untreated mice. For example, in mice treated for four weeks, the correlation coefficient between CFU and 16S rRNA was 0.31 (P-value=0.01) (**Fig 1d**). The R² was 0.10, indicating Only 10% of the variation in CFU was explained by 16S rRNA change alone. Correlations for other treatment durations are shown in Table S7. All regimens decreased CFU to a substantially greater degree than 16S rRNA. Table 2 summarizes the gap between reduction in CFU and 16S rRNA for 9 regimens that have recently been evaluated or are being evaluated in human trials.^27–29^ For example, 4-week treatment with HRZE reduced CFU and 16S rRNA by 2.5 and 0.9 log_10_, respectively, producing a gap of 1.6 log_10_ that corresponds to a 39-times greater reduction in CFU than in 16S rRNA. For bedaquiline (B), pretomanid (Pa), moxifloxacin (M), pyrazinamide (Z) (**BPaMZ**), the gap was more extreme with a reduction in CFU and 16S rRNA by 7.0 and 1.3 log_10_, respectively, producing a gap of 5.7 log_10_ that corresponds to indicating a 527,000-times greater reduction in CFU than in 16S rRNA. Pairwise comparisons between the regimens for both CFU and 16S rRNA are shown in Table S8. The rank order of regimen effects differed depending on whether the outcome was quantified by CFU or 16S rRNA in the regimens as well. For example, CFU ranked BPaMZ as the most potent regimen, whereas it had the fifth greatest effect on 16S rRNA. Similarly, B, Pa, linezolid (L) (**BPaL**) had the fourth greatest effect on CFU but the least effect on 16S rRNA. In contrast, B, Pa, quabodepistat (O), sutezolid (S) (BPaOS) ranked fifth for CFU but showed the greatest effect on 16S rRNA **(Fig. 1g).**

**Table 2.**
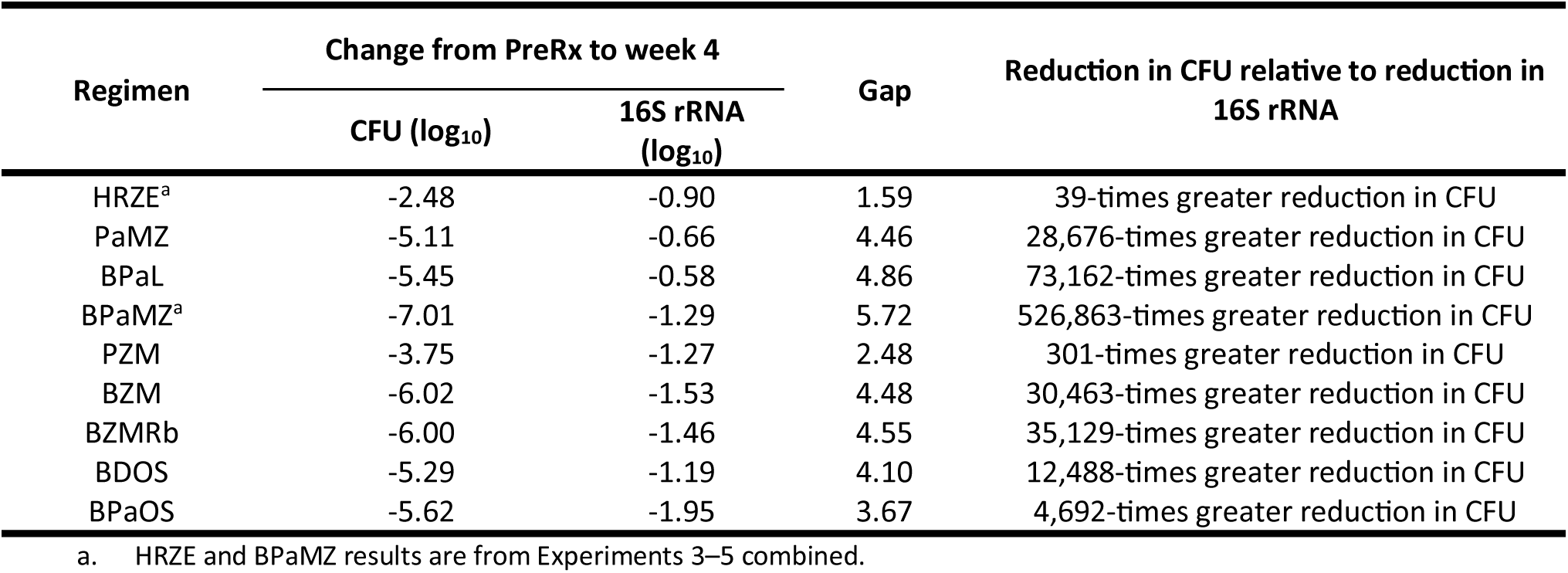
Mean Log_10_ change in lung CFU and 16S rRNA relative to pre-treatment control after four weeks of treatment for selected drug regimens in the BALB/c high-dose aerosol model.

#### Percentage of the gap attributable to residual 16S rRNA from dead Mtb depends on 16S rRNA half-life and time

In experiment 6a, we exposed *Mtb* Erdman strain to HRZE *in vitro* and identified a gap between CFU and 16S rRNA over time **(Fig 2a).** The decline in CFU was biphasic on a log scale whereas the decline in 16S rRNA was nearly exponential. By day 7, CFU decreased 270-times more than 16S rRNA. Using these *in vitro* results, we explored the percentage of the gap that might be attributable to residual 16S rRNA from dead *Mtb* and how this may change over time while assuming a range of values of the half-life times of 16S rRNA from dead bacteria. We began with the simplifying (but likely erroneous) assumption that VBNC_SA_ *Mtb* do not exist and that 16S rRNA comes only from live or dead bacteria (**Fig 2b**). Under this assumption, all viable bacteria are recoverable on agar plates, meaning any decrease in CFU is synonymous with killing. In our novel “Live-Dead” (**LD**) mathematical model (**Fig 2b** and **eqns. (2)-(4)**), antibiotics kill at rate δ_*B*_ and 16S rRNA decays at a rate δ_*S*_, and drug treatment does not change the number of 16S rRNA copies per viable (or dead) bacteria. We modeled expected 16S rRNA counts assuming a range of half-life times for 16S rRNA from dead bacteria. For example, Figure 2c shows the model-predicted 16S rRNA copies if the half-life of 16S rRNA from dead *Mtb* were assumed to be 18 or 24 hours. We then calculated the fraction of gap between CFU and 16S rRNA that is explicable based only on decay of 16S rRNA from dead *Mtb* (**eqn. (6)** and **Fig 2d**). At one theoretical extreme, if the half-life of 16S rRNA were zero (*i.e.,* 16S rRNA disappeared instantaneously when bacteria die, δ_*S*_ → ∞), there would be by definition no residual 16S rRNA from dead *Mtb* and, 0% of the gap would be attributable to residual 16S rRNA from dead *Mtb*. At the opposite extreme, if the half-life of 16S rRNA from dead *Mtb* were assumed to be very long (*e.g.,* 64 hours), 100% of gap could be explained by residual 16S rRNA from dead *Mtb* at most time points (**Fig 2d**). In general, the longer the half-life, the greater the percentage of the gap that would be explained by residual 16S rRNA from dead *Mtb*. Importantly, except for cases when the assumed half-life was longer than the best fit half-life, the percentage of the gap that could be explained by residual 16S rRNA from dead *Mtb* decreased over time (**Fig 2d**). We then fit the LD model to data from mice treated with HRZE for up to 54 days (**Fig S1a-b**). Murine results were similar to *in vitro* in two key respects: (1) the half-life of 16S rRNA from dead *Mtb* would have to be quite long (*i.e.,* >220 hours) to account for most of the gap, and (2) the percentage of the gap that can be explained by residual 16S rRNA generally decreases over time.

**Fig 2.**
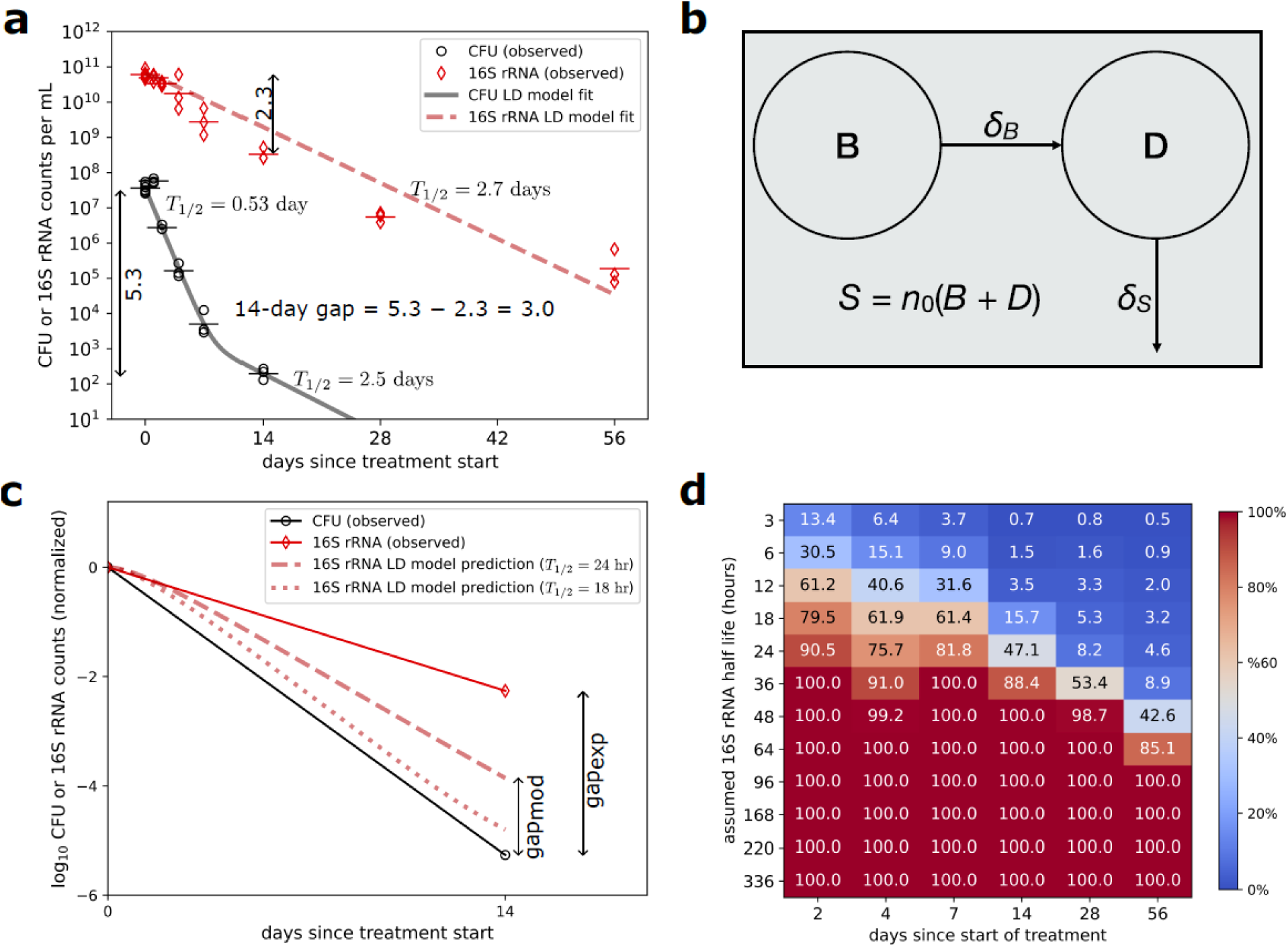
Short half-life time of 16S rRNA from dead bacteria is inconsistent with the gap between CFU and 16S rRNA counts observed during HRZE treatment of Mtb in vitro. We performed experiments by treating in vitro cultures of H37Rv strain of Mtb with HRZE and followed change in platable CFU and 16S rRNA counts in the cultures. **a.** Change in the CFU and 16S rRNA during HRZE treatment. Individual points represent median values at each time point and lines are predictions from a mathematical model assuming division of 16S rRNA pool into live and dead bacteria (see **eqns (2)-(4)**). Note that CFU become undetectable at day 28 and 56. We show the calculation of the gap 14 days after treatment start (gap=3 logs). Best fit parameters are: B_1_(0) = 3.63 x 10^7^, B_2_(0) = 9.41 x 10^3^, D(0) = 0 (assumed), δ_*B*_ = 1.31/day (half-life 0.53 days), δ_*B*_ = _2_ 0.28/day (half-life 2.5 days), δ_*S*_ = 0.26/day (half-life 2.7 days), *n*_0_ = *B*(0)/*S*(0) = 1650 (required by assuming D(0) = 0). **b.** Schematic of the “Live-Dead” (**LD**) mathematical model of Mtb dynamics during treatment. In the model, bacteria die at a rate δ_*B*_and dead bacteria are eliminated/degrade at rate δ_*S*_. The total number of 16S rRNA in the population is *S* = *n*_0_(*B* + *D*) where *n*_0_ is the number of 16S rRNA per cell. **c**. Schematic representation of how the difference in gap was calculated; this includes gap observed in experiments (solid lines) and gap predicted by the matthematical model (dashed line, **eqn. (6)**) assuming that half-life of 16S rRNA in dead bacteria is *T*_1/2_ = 1 day (or δ_*S*_ = 0.69/day) **d.** Percentage gap between the CFU and 16S rRNA markers explained by residual (“leftover”) 16S rRNA assuming different hypothetical half-lives of the 16S rRNA. Values indicate the percentage of the observed gap attributable to remaining 16S rRNA from dead bacteria (**eqn. (6)**). See **Fig S1** for similar calcuations for Mtb dynamics in vivo.

To estimate the half-life of 16S rRNA from dead *Mtb in vitro,* we conducted an *in vitro* experiment in which *Mtb* is known to be killed rapidly. Specifically, in experiment 6b, we exposed *Mtb* Erdman to a lethal concentration (200mM) of hydrogen peroxide (H_2_O_2_). After 24 hours, CFU was reduced from 7.6 log_10_ to undetectable. Nonetheless, 16S rRNA remained quantifiable until day 7. We calculated half-life separately in the first 24 hours after H_2_O_2_ exposure and in the 4 to 7-day interval and identified a 16S rRNA half-life from dead *Mtb* of 5.2 and 15.9 hours respectively. Application of these half-life values in the live-dead model above to HRZE-treated cultures would predict that on day four the percentage of the gap explained by residual 16S rRNA from dead *Mtb* is 12.5% (with 5.2 hour half-life) or 55.5% (with 15.9 hour half-life).

#### Gap between CFU and 16S rRNA after treatment discontinuation

We next examined change in CFU and 16S rRNA during a drug-free recovery period after treatment discontinuation. In Experiment 7, mice were treated for 2 or 4 weeks with HRZE. These durations were intentionally non-curative since it is known that BALB/c infected via high-dose aerosol require ∼ 19-21 weeks to achieve sustained cure in 95% of mice.^7^ When HRZE was stopped after only two weeks, CFU stabilized and then gradually rose during a 4-week drug-free holiday to a CFU count 1.8-times higher than at the end of treatment (**Fig 3a**, **Table 3**). By contrast, in the same mice, 16S rRNA continued to decline during a 4-week recovery phase to a 16S rRNA count 2.6-times lower than at the end of treatment (**Fig 3b**, **Table 3**). When HRZE was stopped after four weeks, CFU returned during a 4-week recovery phase to the end of treatment CFU value (**Fig 3c**, **Table 3).** By contrast, in the same mice, 16S rRNA decreased steadily during the 4-week recovery phase to a 4.7-times lower than at the end of treatment (**Fig 3d**, **Table 3**).

**Fig 3.**
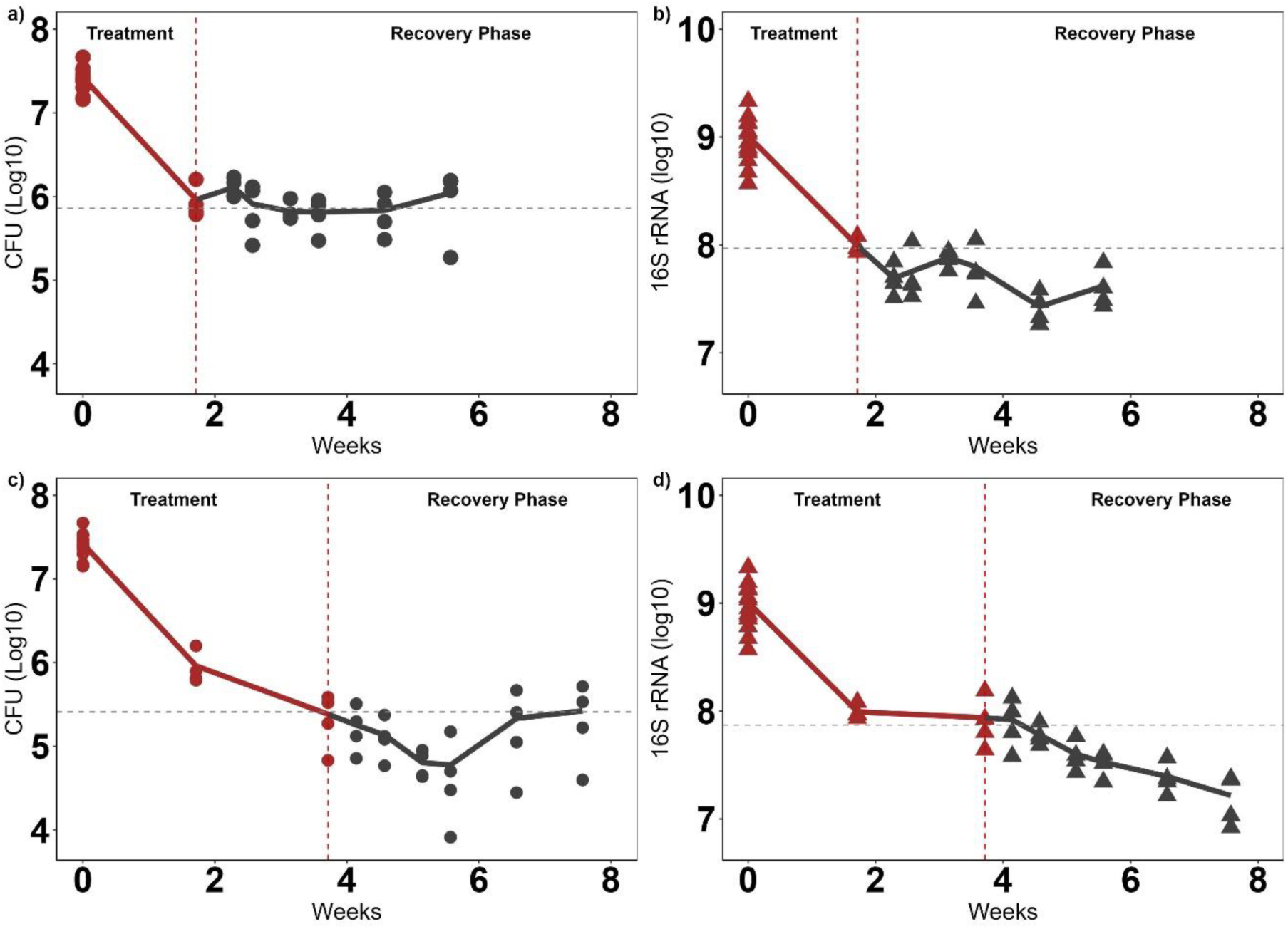
CFU and 16S rRNA when HRZE is stopped after short, non-curative treatment. **a.** CFU during 2-week HRZE treatment followed by a 4-week treatment-free recovery phase in Experiment 6. For all figures in this panel, red symbols represent lung values from individual mice sacrificed at pre-treatment baseline or during treatment. Black symbols represent lung values from individual mice sacrificed 4 to 28 days after final HRZE dose. Lines connect median values. The vertical dotted line marks the end of treatment. The horizonal dotted line marks the end of treatment value. **b**. 16S rRNA during the same treatment time course and recovery described in a. **c**. CFU during 4-week HRZE treatment followed by a 4-week recovery. **d**. 16S rRNA during the same treatment time course and recovery described in c.

**Table 3.** Median log₁₀ CFU, 16S rRNA, and corresponding changes from pre-treatment baseline, including the CFU–16S Gap, in BALB/c mice infected via high-dose aerosol and treated with HRZE for 2 or 4 weeks followed by a 4-week treatment-free recovery period.

| Time point | CFU<br>(log<sub>10</sub>) | 16S rRNA<br>(log<sub>10</sub>) | Change from PreRx (log<sub>10</sub>) |  | Gap |
| --- | --- | --- | --- | --- | --- |
|  |  |  | CFU | 16S rRNA |  |
| Pre-treatment baseline | 7.41 | 9.43 |  |  |  |
| Upon completion of 2-week treatment | 5.86 | 8.45 | -1.55 | -0.98 | 0.61 |
| Upon completion of 2-week treatment<br>followed by 2-week recovery | 5.84 | 8.22 | -1.57 | -1.21 | 0.48 |
| Upon completion of 2-week treatment<br>followed by a 4-week recovery | 6.12 | 8.02 | -1.29 | -1.40 | 0.06 |
| Upon completion of 4-week treatment | 5.39 | 8.34 | -2.02 | -1.08 | 0.93 |
| Upon completion of 4-week treatment<br>followed by a 4-week recovery | 5.38 | 7.67 | -2.03 | -1.75 | 0.23 |
| Upon completion of 8-week treatment | 2.44 | 7.72 | -4.98 | -1.71 | 3.15 |

Second, we examined change in CFU and 16S rRNA during a 12-week drug-free recovery period after treatment with various regimens in several BALB/c RMM studies (Experiments 3-5). Companion groups of mice were sacrificed upon completion of various treatment durations and 12 weeks after the end of treatment (**Table 4**). CFU counts typically rose during the 12-week holiday after sub-curative treatment. By contrast, during the same period, 16S rRNA counts typically fell. For example, following 4-week treatment with BPaL followed by a 12-week drug holiday, CFU count rose 87-times (from 1.9 log at end of treatment to 3.8 log 12 weeks later). By contrast, the 16S rRNA fell 46-times during the same period (from 8.5 log at end of treatment to 6.9 log 12 weeks later) (**Table 4**).

**Table 4.** Change in log_10_ CFU and 16S rRNA counts between end of treatment and relapse assessment 12 weeks after treatment completion. Only treatment arms in which at least one third of mice had a quantifiable CFU result at the end of treatment are shown because median CFU could not be calculated when most samples are below the culture LOD.

| Regimen | Duration (weeks) | Median CFU (log<sub>10</sub>) |  | Change in CFU (log<sub>10</sub>) | Median 16S rRNA (log<sub>10</sub>) |  | Change in 16S rRNA (log<sub>10</sub>) |
| --- | --- | --- | --- | --- | --- | --- | --- |
|  |  | EOT | EOT +12WKS |  | EOT | EOT +12WKS |  |
| BPaL | 4 | 1.91 | 3.85 | 1.94 | 8.54 | 6.88 | -1.66 |
|  | 8 | 0.36 | 1.11 | 0.75 | 7.50 | 6.38 | -1.12 |
| BPaMZ | 2 | 3.75 | 3.40 | -0.35 | 8.53 | 6.55 | -1.98 |
|  | 4 | 0.21 | 2.73 | 2.52 | 8.06 | 6.10 | -1.95 |
| HRZE | 14 | 0.51 | 3.45 | 2.94 | 7.31 | 6.78 | -0.53 |
| BDOS | 4 | 1.96 | 3.91 | 1.95 | 8.10 | 6.30 | -1.80 |
|  | 6 | 0.19 | 3.40 | 3.20 | 7.73 | 6.26 | -1.47 |
| BPaOS | 4 | 1.83 | 3.96 | 2.13 | 7.36 | 6.44 | -0.91 |
|  | 6 | 0.49 | 2.08 | 1.58 | 7.75 | 6.65 | -1.10 |

**Table 5.** Output from meta-regression modeling of T_95_ as a function of CFU (Model 1), 16S rRNA (Model 2), or both CFU and 16S rRNA (Model 3). The likelihood ratio test (LRT) full-model comparison evaluates whether the full model (Model 3) provides a significantly better fit than each reduced model (Models 1 and 2). The LRT null comparison evaluates whether each model differs significantly from the null model containing no predictors.

| Model | Predictor Variables | $R^2$ | AIC | RMSE | LRT P- value (full model comparison) | LRT P-value (null comparison) |
| --- | --- | --- | --- | --- | --- | --- |
| 1 | CFU (Baseline + average 28-day change) | 0.90 | 52.95 | 2.05 | 0.019 | <0.001 |
| 2 | 16S rRNA (Baseline + average 28-day change) | 0.24 | 71.90 | 4.40 | <0.001 | 0.228 |
| 3 | CFU (Baseline + average 28-day change) + 16S rRNA (Baseline + average 28-day change) | 0.96 | 49.00 | 1.56 | - | <0.001 |

#### CFU and 16S rRNA as predictors of relapse in the RMM

We evaluated the association of CFU and 16S rRNA with microbiologic relapse in three RMM studies conducted in two different laboratories (experiments 3-5). The ability of regimens to prevent relapse (*i.e.,* durable cure) was quantified in terms of T_95_ for 9 unique three- or four-drug regimens (**Fig. 4a and Table 3**). Because HRZE and BPaMZ were used as controls in several trials, these regimens have several independent estimates, resulting in 11 individual arms. The shortest and longest T_95_ were 5.06 weeks for BPaMZ and 20.8 weeks for HRZE.

**Fig 4.**
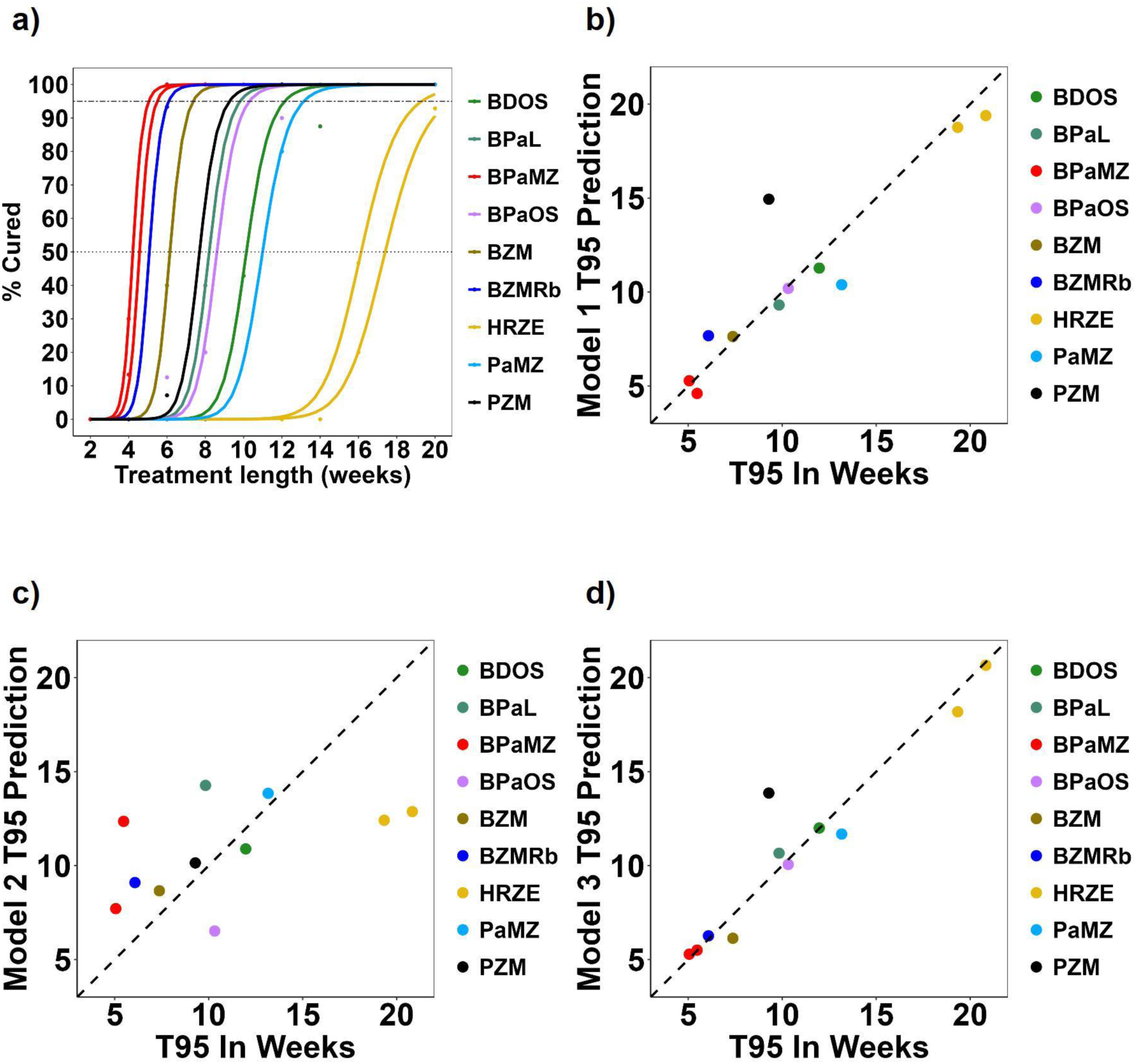
T_95_ and model outputs. **a.** Estimated cure rate and the duration of treatent for each regimen evaluated in three BALB/c relapsing mouse model experiments used to estimate T_95_ (upper dotted horizontal line). **b-d,** Correlation between the observed T_95_ and predicted T_95_ for Models 1-3, respectively.

Using multivariable meta-regression, we first modeled T_95_ as a function of CFU alone (Model 1: Pretreatment CFU + week 4 CFU change). The Model 1 R^2^ was 0.90 and predicted T_95_ values were strongly associated with the observed T_95_ (correlation coefficient 0.91, *P*=8.3×10^-05^ (**Fig 4b**). We then modeled T_95_ as a function of 16S rRNA alone (Model 2: Pretreatment 16S rRNA + week 4 16S rRNA change). The Model 2 R^2^ was 0.24 and predicted T_95_ values were weakly and non-significantly associated with observed T_95_ (correlation coefficient 0.49, *P*=0.13) (**Fig 4c**). Finally, we modeled T_95_ as a function of both CFU and 16S rRNA (Model 3: Pretreatment CFU and 16S rRNA + 4-week CFU and 16S rRNA change). The Model 3 R^2^ was 0.96 and predicted T_95_ values were strongly associated with observed T_95_ (correlation coefficient 0.95, *P*=5.7 x10^-06^) (**Fig 4d**). Use of both CFU and 16S rRNA in Model 3 slightly improved upon Model 1 which used CFU alone (likelihood ratio test *P-*value 0.019).

## DISCUSSION

To address an important uncertainty about PD markers used in murine TB drug evaluation, we evaluated discordance between CFU and 16S rRNA in the BALB/c mouse model that is a reference standard in preclinical development. We found that 16S rRNA was an excellent surrogate for CFU in untreated mice, but a large gap emerged between 16S rRNA and CFU values during drug treatment. Combination drug regimens typically reduced CFU thousands of times more than 16S rRNA. Our modeling of *in vitro* results showed that, unless the half-life of 16S rRNA were assumed to be ≥96 hours, the fraction of the gap potentially attributable to delayed clearance of residual 16S rRNA from dead *Mtb* decreased over time. This suggests that excess 16S rRNA observed at later time points might originate from a VBNC_SA_ population. When treatment was discontinued in mice, we found that 16S rRNA decreased over time even as CFU rose, suggesting either very slow decay of 16S rRNA from dead *Mtb* or progressive elimination of a VBNC_SA_ population. Finally, change in 16S rRNA alone was not significantly associated with the capacity of regimens to rapidly cure TB (quantified as T_95_). However, change in 16S rRNA did significantly improve accuracy of prediction of T_95_ when included in a combined CFU and 16S rRNA model. In the aggregate, these findings help to identify and partially constrain the potential causes of discordance between CFU and 16S rRNA results in murine treatment models, enhancing interpretation of these PD markers.

We analyzed results from diverse murine and *in vitro* experimental designs to address the puzzling and currently unexplained gap between CFU and 16S rRNA results. Inferences drawn from each experiment are discussed sequentially below.

Our results in the drug-naive “chronic” BALB/c mouse model (Experiment 1) reinforced existing evidence^18^ that, in the absence of drug treatment, 16S rRNA and CFU provide similar information about *Mtb* burden (R-squared = 0.92). Log 16S rRNA was notably related to log CFU with a slope of 0.83 rather than 1, suggesting that the quantity of 16S rRNA per bacillus could decrease as the onset of adaptive immunity results in a slower growing *Mtb* phenotype.

Our results in BALB/c mice treated with individual drugs (Experiment 2) expanded upon previous evidence that drugs affect CFU and 16S rRNA divergently in mice. As reported by Evangelopoulos *et.al.*^18^, we found that four weeks of rifampin 30mg/kg decreased CFU 67-times more than 16S rRNA. Similarly, bedaquiline and streptomycin also decreased CFU substantially more than 16S rRNA. Conversely, a novel observation was that isoniazid, pyrazinamide and ethambutol decreased CFU less than 16S rRNA. Collectively, these monotherapy results confirm that 16S rRNA is not a straightforward substitute for CFU during treatment with individual drugs in BALB/c mice. The relationship between CFU and 16S rRNA is variable and appears to depend on drug mechanism and duration of treatment.

Our results in BALB/c mice treated with drug combinations (experiments 3-5) showed that all regimens reduced CFU to a greater degree than 16S rRNA. There was substantial regimen-to-regimen variation in the magnitude of the gap between CFU and 16S rRNA. For example, four weeks of BPaMZ reduced CFU around 527,000 times more than 16S rRNA, whereas HRZE reduced CFU only ∼39 times more than 16S rRNA. Neither the monotherapy nor combination results alone indicate whether the gap signals a shift of *Mtb* to a VBMC_SA_ status or indicates persistent detection of residual 16S rRNA from dead *Mtb*.

Our *in vitro* experiment with HRZE replicated the gap between CFU and 16S rRNA observed *in vivo.* Mathematical modeling of these *in vitro* or *in vivo* results yielded two key findings. First, the percentage of this gap that could be explained by residual 16S rRNA from dead *Mtb* depended on how long 16S rRNA is assumed to persist after *Mtb* was killed. The longer the half-life, the more of the gap could be attributed to residual 16S rRNA from dead *Mtb.* Second, irrespective of the half-life assumption, the percentage of the gap that could be attributed to residual 16S rRNA from dead *Mtb* decreased over time. This suggests that over time, an increasing share of the “excess” 16S rRNA not explained by CFU may be due to VBNC_SA_. Our *in vitro* experiment with H_2_O_2_ had two key findings. First, it showed that 16S rRNA may remain detectable after *Mtb* death. Second, it identified a range of plausible half-life estimates for 16S rRNA from dead *Mtb in vitro.* Assuming the longer half-life estimate (15.9 hours or 0.7 days) is accurate, residual 16S rRNA from dead *Mtb* would account for about half the gap at day 4 but only ∼8% at day 14 in our *in vitro* experiments. With shorter half-life assumptions, the proportion of the gap explained by residual 16S rRNA from dead *Mtb* became progressively smaller. This discovery of “excess” 16S rRNA not explained by CFU or dead *Mtb* based on the *in vitro* determined half-life of 16S rRNA from dead bacteria implies the presence of VBNC_SA_.

Evaluation of BALB/c mice when treatment was discontinued prior to cure showed that CFU “rebounded” during a drug-free holiday yet 16S rRNA continued to decrease. We see two possible explanations. First, it is conceivable that there is a large pool of 16S rRNA from already-killed *Mtb* that continues to slowly decay even as viable *Mtb* resume replication. However, for this to be the only explanation, 16S rRNA from dead *Mtb* would have to have a very long half-life *in vivo* (around 7-10 days). Second, it is possible that antibiotic-damaged *Mtb* become non-culturable as they progress towards death. This would suggest that the VBNC_SA_ may be a pre-terminal stage rather than a stable state that *Mtb* can sustain indefinitely. Future studies will need to be designed to discriminate between these alternatives.

Since the above analyses indicated that CFU and 16S rRNA likely measure correlated but different phenomena, we were particularly interested in the association of CFU or 16S rRNA with relapse in the conventional BALB/c model. For the nine unique regimens tested, change in CFU alone was robustly associated with T_95_ but change in 16S rRNA alone was not. However, addition of 16S rRNA results to the CFU model did significantly enhance the association with T_95_, increasing R^2^ from 0.90 to 0.96. This suggests that 16S rRNA does provide information distinct from CFU, even if 16S rRNA alone is a relatively poor predictor of T_95_.

This study has several limitations. First, we used a single mouse model (BALB/c) that is the contemporary standard in preclinical drug evaluation. Future work could evaluate alternative models such as the C3HeB/FeJ mouse. Our murine results should not be extrapolated to the use of *Mtb* 16S rRNA for measurement of treatment responses in sputum which is being evaluated in ongoing human trials. Second, our experiments did not perform Most Probable Number (MPN) testing, a culture-based dilution assay that identifies “differentially culturable” *Mtb* that fail to grow on agar and are therefore not quantified by CFU.^30^ Third, because only nine unique regimens were available for modeling the association between CFU or 16S rRNA and relapse (*i.e.,* T_95_), we consider their associations with relapse as exploratory. Fourth, our basic two-compartment model had several simplifying assumptions. Most importantly, it included only culturable and dead *Mtb* and required some simplification to account for the biexponential decline in CFU. Future work will evaluate more complex mathematical models allowing for different sources of 16S rRNA (e.g., from dead and VBNC_SA_ bacteria) and allowing the number of rRNA copies per bacterium to change between populations and over time. Fifth, a key parameter of our model (*in vivo* half-life of 16S rRNA from dead *Mtb*) is highly uncertain. We therefore illustrated the full range of possible scenarios, ranging from a very short to a very long half-life time. Finally, for the purposes of clarity, we have framed our analysis based on two dueling hypotheses (shift to non-culturable state versus slow clearance of residual 16S rRNA). However, these two phenomena are not mutually exclusive. It is notoriously difficult to “prove the negative” and demonstrate the absence of microbial viability.^31^ Nonetheless, we believe the preponderance of our observations point to both slow clearance of residual 16S rRNA from dead *Mtb* and the presence of a VBNC_SA_ population as contributors to the observed gap. The timepoint at which the PD marker is measured likely determines which of these two explanations is the predominant cause. Early timepoints suggest that the majority of the gap arises from slow clearance of dead bacteria, while later timepoints point to the presence of a VBNC_SA_ population.

## CONCLUSION

During treatment with regimens both *in vitro* and *in vivo*, a gap develops in which there is a large excess of 16S rRNA relative to CFU. In the aggregate, our findings suggest that the gap may be due to a combination of slow decay of residual 16S rRNA from already-killed *Mtb* and development of VBNC_SA_ population, with the latter becoming progressively more dominant over time. However, our observations that 16S rRNA continued to decrease after treatment cessation even as CFU increased and the poor association of change in 16S rRNA with relapse suggests that the VBNC_SA_ may be damaged to the point of being irrecoverable and thus may be equivalent to dead *Mtb*. Broadly, this work highlights the complexity of establishing exactly what PD markers are measuring and the need for deeper understanding and continued innovation to improve measurement of drug effects.

## Abbreviations

TB: tuberculosis
Mtb: Mycobacterium tuberculosis
LD: live-dead.

## Acknowledgements

We gratefully acknowledge insightful comments from Dr Patrick Phillips on our draft manuscript.

## Author contributions

**Samuel T. Tabor**: Conceptualization; Methodology; Investigation; Data curation; Formal analysis; Visualization; Interpretation; Writing – original draft.

**Allan D. Friesen**: Methodology (mathematical modeling); Formal analysis; Visualization; Interpretation; Writing – review & editing.

**Matthew J. Reichlen**: Methodology; Investigation; Technical oversight; Writing – review & editing.

**Christian Dide-Agossou**: Data curation; Formal analysis.

**Max McGrath** and **Ryan Peterson**: Formal analysis (meta-multivariable regression); Interpretation.

**Vitaly V. Ganusov**: Methodology (mathematical modeling); Interpretation; Critical review of analytical approach; Writing – review & editing; Funding acquisition.

**Gregory T. Robertson**: Conceptualization; Methodology; Supervision; Interpretation; Writing – review & editing; Funding acquisition.

**Martin I. Voskuil**: Conceptualization; Methodology; Supervision (experiments); Interpretation; Writing – review & editing; Funding acquisition.

**Nicholas D. Walter**: Conceptualization; Methodology; Supervision (experiments and analytical work); Interpretation; Writing – review & editing; Writing – original draft support; Funding acquisition.

## Data and materials availability

All Data and Code are available at the following link: https://github.com/SamuelTaborCU/Mtb-16S-rRNA-vs-CFU and https://github.com/allanfriesen/CFU16SrRNAgap

