## Supplemental information for "Mind the gap: Understanding discordance between culture- and a non-culture-based measure of bacterial burden in murine tuberculosis treatment models"

### **Supplemental Methods**

#### **Enumeration of colony forming units (CFU)**

#### **From lungs**

The number of viable organisms was determined by serial dilutions of homogenates (Precellys Evolution, Bertin) prepared in phosphate buffered saline plus 10% (w/v) bovine serum albumin from indicated lung lobes and plating on 7H11-OADC agar plates containing 0.4% (w/v) activated charcoal to prevent drug carry-over. Colonies were enumerated after at least 21 days of incubation at 37°C. For relapse assessments, tissues were homogenized in PBS and plated in their entirety on 7H11-OADC agar plates without activated charcoal.

#### **From *In vitro***

The number of viable organisms was determined by serial dilutions of *M. tuberculosis* cultures prepared in complete 7H9 medium. Twenty microliters of undiluted culture was placed into the first well of a 96-well microtiter plate, and six additional 10-fold serial dilutions (10⁻¹ to 10⁻⁶) were prepared by transferring 20 µL into 180 µL of medium. 4 μL from the neat culture and each dilution was spotted in triplicate onto DTA agar plate with 1.5% Bacto agar, 10% OADC supplement and 0.4% activated charcoal. Colonies were enumerated after at least 21 days of incubation at 37 °C.

#### **RNA extraction**

Lung tissues from murine samples were beadbeaten in 2 ml Trizol using the CKmix50 lysis kit for 1×16 seconds at 7,200 rpm followed by 3×30 seconds at 6,500 rpm with 5-minute rests on wet ice between cycles using the CK01 lysis kit (Bertin). Cellular debris was separated from the cell lysate by centrifugation for 1 minute at 21,000×g. Trizol lysates were transferred to a new tube containing heavy phase lock gel and 300 μl chloroform, mixed vigorously for 15 seconds, then incubated at room temperature for 2 minutes with occasional mixing followed by centrifugation at 21,000×g for 10 minutes. The aqueous phase was removed to a separate tube for RNA extraction and mixed with 270 μl of isopropanol. 265 μl of high salt solution (0.8 M sodium citrate, 1.2 M sodium chloride) was added and mixed by inversion. RNA was precipitated overnight at 4°C and then pelleted by centrifugation at 4°C, 21,000×g for 10 minutes. The RNA pellet was air dried for 10 minutes at room temperature, resuspended in 80 µl water, and reconstituted for one hour on ice. 2 μl DNase I and 10 μl DNase I buffer (Promega) and 2.5 μl Ribolock RNase inhibitor were added to the resuspended pellet and incubated for 30 minutes at 37°C. RNA was then purified with the Maxwell RSC simplyRNA tissue kit using the Maxwell RSC instrument (Promega) following the manufacturer’s instructions with the following modifications: an additional DNase treatment was performed with Promega RQ1 DNase prior to instrument use and additional kit DNase was added at twice the recommended amount for instrument use. For every extraction batch, a negative extraction control was included to test cross-contamination between samples. RNA extraction efficiency was tracked by spiking in a commercially available external RNA control mix (ERCC) (Ambion, # 4456740) immediately prior to extraction. The RNA extraction efficiency was calculated as a percentage of RNA that was retained during the extraction process. To track RNA retention percentage, an equal volume of the ERCC mix from the same stock was diluted to match the final RNA elution volume.

#### **Evaluation of the adjusting for RNA retention for the calculation of rRNA burden**

We evaluated whether adjusting 16S rRNA burdens for RNA retention (i.e., extraction efficiency) improves precision. For each extraction, we estimated RNA-retention percentage using ERCC spike-ins. Using multiple technical and biological replicates, we first quantified concordance between adjusted and unadjusted burdens, then compared %CV across replicate sets. Adjustment did not improve precision: adjusted and unadjusted burdens were highly concordant (Pearson r ≈ 0.90), and %CVs were typically larger after adjustment in both biological and technical replicates. A one-tailed paired t-test on paired %CVs, testing whether adjustment reduces variability, provided no evidence of improvement (t = 1.94, p = 0.973).

#### **Quantification of rRNA burden via RT-qPCR**

After extraction, RNA was reverse transcribed using the SuperScript VILO cDNA Synthesis Kit (Invitrogen), with incubation at 25 °C for 10 min, 50 °C for 10 min, 85 °C for 5 min, and a final hold at 4 °C. The resulting cDNA was diluted 1:10. Two microliters of diluted cDNA were used as input for RT-qPCR (TaqMan assay) for absolute quantification of 16S rRNA. All samples were run in triplicate, and median values were used for analysis. Primer and probe sequences are provided in Table S10. RT-qPCR was performed under the following thermocycling conditions: initial denaturation at 95 °C for 10 min; 40 cycles of 94 °C for 30 s and 60 °C for 1 min; followed by a final hold at 4 °C. *M. tuberculosis* rRNA burden was expressed as the absolute number of 16S rRNA transcripts, adjusted for input RNA volume and dilution factors.

#### **Description of the Emax model**

The sigmoidal E_max_ model was tested according to:

$P_{cure} =\left( 1-P_{relapse} \right)= E_{0}+$ $\frac{E_{max} \times T^{\gamma}}{T_{50}^{\gamma}+T^{\gamma}}$	(1)

In this model, P_cure_ is the probability of cure defined as a negative solid culture 3 months post-treatment. The independent variable is treatment length. E_0_ is the expected response when exposure is zero (*i.e.,* basal effect). E_max_ is the maximal achievable probability of cure. The E_0_ and E_max_ parameters were constrained to 0 and 1, respectively. γ is a shape parameter controlling the steepness of the curve produced by the E_max_ equation. T_50_ is the treatment length at which half the E_max_ is achieved. From the T_50_ estimates we derived T_95_ of the combination regimens according to the formula:

$T_{95}= T_{50}\times\left( \frac{95}{100 -95} \right)^{1/\gamma}$	(2)

#### **Description of BALB/c mouse experiments**

*Experiment 1: BALB/c mouse low dose “chronic” aerosol infection model*

Seven to nine -week-old female pathogen-free BALB/c mice (Jackson Laboratories) were exposed to *low dose “chronic” aerosol* of *M. tuberculosis* Erdman using the Glas-Col Inhalation Exposure System to achieve deposition of ~2.4 log10 CFU in the lungs of each mouse one day post-infection. Five mice were sacrificed on days 1, 7, 11, 19, 28, and 56 post-infection for sample collection.

*Experiment 2: BALB/c mouse high dose aerosol infection model*

Six to 8-week-old female pathogen-free BALB/c mice (Jackson Laboratories) were exposed to high-dose aerosol of *M. tuberculosis* Erdman from broth culture (OD_600_ ~ 0.8) to achieve deposition of ~3.8 log10 CFU in the lungs of each mouse.^1,2^ Treatment was initiated on day 11 post aerosol and continued for 4 weeks.^3^ Groups of 6 mice were individually euthanized by CO_2_ narcosis on day 11, prior to treatment initiation, and on the last day of treatment, to determine the bacterial loads in lungs. The left and lower right lung lobes (inferior and post-caval lobes) were used for bacterial enumeration. Upper right lung lobes (superior and middle lobes) were flash frozen in liquid nitrogen prior to RNA extraction.

*Experiments 3, 4 and 5: BALB/c relapsing mouse models*

Six to 8-week-old female pathogen-free BALB/c mice (Jackson Laboratories) were exposed to high-dose aerosol of *M. tuberculosis* from broth culture (OD_600_ ~0.8) to achieve deposition of ~ 4.3 log_10_ CFU in the lungs of each mouse.^1,2^ Treatment was initiated on post-infection day 15 (Experiment 2 at JHU) or day 11 (Experiment 3 and 5 at CSU) and continued for up to 20 weeks. Groups of 3 to 6 mice, as indicated, were individually euthanized by CO_2_ narcosis prior to treatment initiation, and one day following the last day of treatment, to determine the bacterial loads in the lungs. Additional groups of 15 mice each from each treatment group were placed on a 12-week drug holiday for evaluation of the conventional microbiological relapse outcome.^1,4–7^ The left and lower right lung lobes (inferior and post-caval lobes) were used for bacterial enumeration. Upper right lung lobes (superior and middle lobes) were flash frozen in liquid nitrogen prior to bead-beating and RNA extraction.

*Experiments 7: BALB/c “rebound”* *post treatment models*

Six- to 8-week-old female pathogen-free BALB/c mice (Jackson Laboratories) were infected using a high-dose aerosol exposure (Glas-Col) with *M. tuberculosis* Erdman, resulting in a mean lung implantation of 4.43 log₁₀ CFU on day 1. On day 11 post-infection, five mice were euthanized by CO₂ narcosis as pre-treatment controls. The remaining mice initiated therapy with HRZE administered by oral gavage five days per week, using established standard doses for all antibiotics (**Table S5**). End-of-treatment mice (N = 4 per treatment duration) were sacrificed on days 12, 26, and 54 after treatment initiation, corresponding to 2, 4, and 8 weeks of HRZE treatment, respectively, one day after the final dose, to assess bacterial burden.

Additional groups of mice (N = 4 per time point) from the 12- and 26-day treatment groups were euthanized on days 5, 7, 11, 14, 21, and 28 after treatment interruption to assess physiologic recovery of *M. tuberculosis* **(Fig. 3a–d**). Lung lobes were dissected such that the left lung and lower right lung lobes (inferior and post-caval) were processed for CFU enumeration, while upper right lung lobes (superior and middle) were flash-frozen in liquid nitrogen prior to RNA extraction, as previously described.

**SUPPLEMENTAL FIGURES & TABLES**

#### **Table S1. Experiment table**

| **Experiment** | **Site** | **Strain** | **Inoculum** | **Type** | **Publication** |
| --- | --- | --- | --- | --- | --- |
| *1* | CSU | Erdman | Low-dose aerosol | Untreated | Unpublished |
| *2* | CSU | Erdman | High-dose aerosol | Single drug exposure | PMID: 34006838^3^,  PMID: 35311519^8^ |
| *3* | CSU | Erdman | High-dose aerosol | RMM | PMID: 34006838^3^,  PMID: 35311519^8^ |
| *4* | JHU | H37Rv | High-dose aerosol | RMM | PMID: 35311519^8^ |
| *5* | CSU | Erdman | High-dose aerosol | RMM | Unpublished |
| *6* | UC-AMC | Erdman |  | In vitro | Unpublished |
| *7* | CSU | Erdman | High-dose aerosol | "Rebound" post treatment | Unpublished |

#### **Table S2. Antibiotic dosage concentrations for mice in Experiment 2**.

Treatments began 11 days post-infection and were administered by oral gavage or subcutaneous (s.c.) injection, five days per week, for the designated treatment duration prior to euthanasia.

| **Drug** | **Dose (mg/kg)** |
| --- | --- |
| Bedaquiline | 5 and 25 |
| Ethambutol | 100 |
| Isoniazid | 25 |
| Pyrazinamide | 150 |
| Rifampin | 10 and 30 |
| Streptomycin (s.c.) | 200 |

#### **Table S3. Antibiotic dosage concentrations for mice in Experiments 3, 4, and 5.**

Treatments began 11/15 days post-infection and were administered by oral gavage, five days per week, for the designated treatment duration prior to euthanasia.

| **Regimen** | **Drug 1** | **Dose (mg/kg)** | **Drug 2** | **Dose (mg/kg)** | **Drug 3** | **Dose (mg/kg)** | **Drug 4** | **Dose (mg/kg)** |
| --- | --- | --- | --- | --- | --- | --- | --- | --- |
| **HRZE** | Isoniazid | 10 | Rifampin | 10 | Pyrazinamide | 150 | Ethambutol | 100 |
| **PaMZ** | Pretomanid | 50 | Moxifloxacin | 100 | Pyrazinamide | 150 |  |  |
| **BPaL** | Bedaquiline | 25 | Pretomanid | 100 | Linezolid | 100 |  |  |
| **BPaMZ** | Bedaquiline | 25 | Pretomanid | 100 | Moxifloxacin | 100 | Pyrazinamide | 150 |
| **PZM** | Rifapentine | 10 | Pyrazinamide | 150 | Moxifloxacin | 100 |  |  |
| **BZM** | Bedaquiline | 25 | Pyrazinamide | 150 | Moxifloxacin | 100 |  |  |
| **BZMRb** | Bedaquiline | 25 | Pyrazinamide | 150 | Moxifloxacin | 100 | Rifabutin | 5 |
| **BDOS** | Bedaquiline | 25 | Delamanid | 6 | Quabodepistat | 9 | Sutezolid | 50 |
| **BPaOS** | Bedaquiline | 25 | Pretomanid | 50 | Quabodepistat | 9 | Sutezolid | 50 |

#### **Table S4. Drug dosage concentrations for Experiment 6 in the *in vitro* study.**

| **Regimen** | **Drug 1** | **Dose** | **Drug 2** | **Dose** | **Drug 3** | **Dose** | **Drug 4** | **Dose** |
| --- | --- | --- | --- | --- | --- | --- | --- | --- |
| HRZE | Isoniazid | 0.5 µg/mL | Rifampin | 0.5 µg/mL | Pyrazinamide | 16 µg/mL | Ethambutol | 20 µg/mL |
| H₂O₂ | H₂O₂ | 200mM |  |  |  |  |  |  |

#### **Table S5. Antibiotic dosage concentrations for mice in Experiment 7**.

Treatments began 11 days post-infection and were administered by oral gavage, five days per week, for the designated treatment duration.

| **Drug** | **Dose (mg/kg)** |
| --- | --- |
| Isoniazid | 10 |
| Rifampin | 10 |
| Pyrazinamide | 150 |
| Ethambutol | 100 |

#### **Table S6. Effect of individual drugs on CFU and 16S rRNA in Experiment 2.**

Log_10_ decrease relative to pretreatment control and pairwise comparison *P*-values between mice treated for 4 weeks with pyrazinamide (PZA), ethambutol (EMB), streptomycin (STR), rifampin (RIF), isoniazid (INH) and bedaquiline (BDQ). Pairwise comparisons between individual drugs were performed using two-sample Wilcoxon tests. Average log_10_ decrease are shown in the top row and left-hand column. *P*-values are shown in black. Statistically significant *P*-values are highlighted in bold.

| **CFU** | | | | | | | | | |
| --- | --- | --- | --- | --- | --- | --- | --- | --- | --- |
|  |  | BDQ25 | BDQ5 | RIF10 | RIF30 | STR | EMB | PZA | INH |
|  | *log_10_ decrease* | *3.8* | *2.6* | *1.1* | *2.9* | *0.5* | *0.1* | *0.1* | *1.2* |
| BDQ25 | *3.8* |  | **0.001** | **0.0009** | **0.0009** | **0.0009** | **0.0009** | **0.0009** | **0.001** |
| BDQ5 | *2.6* |  |  | **0.0003** | **0.01** | **0.0003** | **0.001** | **0.0003** | **0.0006** |
| RIF10 | *1.1* |  |  |  | **0.0002** | **0.02** | **0.0009** | **0.0002** | 0.5 |
| RIF30 | *2.9* |  |  |  |  | **0.0002** | **0.0009** | **0.0002** | **0.0003** |
| STR | *0.5* |  |  |  |  |  | **0.0009** | **0.0002** | **0.0003** |
| EMB | *0.1* |  |  |  |  |  |  | 0.8 | **0.001** |
| PZA | *0.1* |  |  |  |  |  |  |  | **0.0003** |

| **16S rRNA** | | | | | | | | | |
| --- | --- | --- | --- | --- | --- | --- | --- | --- | --- |
|  |  | BDQ25 | BDQ5 | RIF10 | RIF30 | STR | EMB | PZA | INH |
|  | *log_10_ decrease* | *1.9* | *1.6* | *0.5* | *1.1* | *0.02* | *0.9* | *0.6* | *1.9* |
| BDQ25 | *1.9* |  | **0.02** | **0.0002** | **0.0002** | **0.0002** | **0.0002** | **0.0002** | 1.0 |
| BDQ5 | *1.6* |  |  | **0.0003** | **0.0006** | **0.0003** | **0.0003** | **0.0003** | 0.07 |
| RIF10 | *0.05* |  |  |  | **0.0002** | **0.0006** | **0.0002** | **0.04** | **0.0003** |
| RIF30 | *1.1* |  |  |  |  | **0.0002** | **0.003** | **0.0006** | **0.0003** |
| STR | *0.02* |  |  |  |  |  | **0.0002** | **0.0002** | **0.0003** |
| EMB | *0.9* |  |  |  |  |  |  | **0.005** | **0.0003** |
| PZA | *1.9* |  |  |  |  |  |  |  | **0.0003** |

#### **Table S7. Correlations between CFU and 16S rRNA for different treatment durations in Experiments 3, 4 and 5.**

| **Treatment Days** | **Corr Coefficient** | **CI_L** | **CI_U** | **p Value** |
| --- | --- | --- | --- | --- |
| **7** | -0.66 | -0.9 | -0.14 | 0.02 |
| **14** | 0.53 | 0.33 | 0.69 | 0.000006 |
| **21** | -0.47 | -0.73 | -0.08 | 0.02 |
| **28** | 0.31 | 0.07 | 0.52 | 0.01 |
| **42** | 0.6 | 0.28 | 0.8 | 0.001 |
| **56** | 0.34 | -0.01 | 0.61 | 0.06 |
| **70** | 0.73 | 0.34 | 0.9 | 0.002 |
| **98** | 0.57 | -0.45 | 0.94 | 0.2 |
| **112** | 0.3 | -0.33 | 0.75 | 0.3 |

#### **Table S8. Effect of combination regimens on CFU and 16S rRNA in Experiments 3, 4 and 5.**

Log_10_ decrease relative to pretreatment control and pairwise comparison *P* values among mice treated for 4 weeks with HRZE, PaMZ, BPaL, BPaMZ, PZM, BZM, BZMRb, BDOS, and BPaOS. Pairwise comparisons between combination regimens were performed using two-sample Wilcoxon tests. Average log_10_ decreases are shown in the top row and left-hand column. P-values are shown in black, and statistically significant P-values are highlighted in bold.

| **CFU** | | | | | | | | | | |
| --- | --- | --- | --- | --- | --- | --- | --- | --- | --- | --- |
|  |  | HRZE | PaMZ | BPaL | BPaMZ | PZM | BZM | BZMRb | BDOS | BPaOS |
|  | *log_10_ decrease* | 2.48 | 5.11 | 5.45 | 7.01 | 3.75 | 6.02 | 6 | 5.29 | 5.62 |
| HRZE | 2.48 |  | **0.0005** | **0.0005** | **0.00002** | **0.003** | **0.001** | **0.001** | **0.001** | **0.003** |
| PaMZ | 5.11 |  |  | 0.2 | **0.001** | **0.01** | **0.04** | **0.01** | 0.4 | 0.2 |
| BPaL | 5.45 |  |  |  | **0.001** | **0.01** | 0.05 | **0.02** | 0.9 | 0.9 |
| BPaMZ | 7.01 |  |  |  |  | **0.004** | **0.003** | **0.004** | **0.002** | **0.01** |
| PZM | 3.75 |  |  |  |  |  | **0.02** | **0.02** | **0.02** | **0.03** |
| BZM | 6.02 |  |  |  |  |  |  | 0.9 | 0.06 | 0.6 |
| BZMRb | 6 |  |  |  |  |  |  |  | **0.03** | 0.3 |
| BDOS | 5.29 |  |  |  |  |  |  |  |  | 0.7 |

| **16S rRNA** | | | | | | | | | | |
| --- | --- | --- | --- | --- | --- | --- | --- | --- | --- | --- |
|  |  | HRZE | PaMZ | BPaL | BPaMZ | PZM | BZM | BZMRb | BDOS | BPaOS |
|  | *log_10_ decrease* | 0.9 | 0.66 | 0.58 | 1.29 | 1.27 | 1.53 | 1.46 | 1.19 | 1.95 |
| HRZE | 0.9 |  | 0.05 | **0.001** | **0.03** | **0.04** | **0.01** | **0.004** | 0.3 | **0.0004** |
| PaMZ | 0.66 |  |  | 0.6 | **0.001** | **0.01** | **0.004** | **0.009** | 0.05 | **0.01** |
| BPaL | 0.58 |  |  |  | **0.0002** | **0.01** | **0.004** | **0.004** | **0.02** | **0.01** |
| BPaMZ | 1.29 |  |  |  |  | 0.8 | 0.3 | 0.6 | 0.7 | 0.1 |
| PZM | 1.27 |  |  |  |  |  | 0.7 | 0.4 | 0.9 | 0.1 |
| BZM | 1.53 |  |  |  |  |  |  | 0.8 | 0.3 | 0.6 |
| BZMRb | 1.46 |  |  |  |  |  |  |  | 0.4 | 0.3 |
| BDOS | 1.19 |  |  |  |  |  |  |  |  | 0.2 |

#### **Table S9 Half-life estimation from in vitro H2O2 exposure.**

| Type | Treatment Days | Half-Life (hours) |
| --- | --- | --- |
| Interval | 1 | 5.2 |
| Interval | 2 | 13.9 |
| Interval | 4 | 12.7 |
| Interval | 7 | 15.9 |
| Overall |  | 12.3 |

#### **Table S10. Primer sequences & information.**

| Primer/Probe name | Sequence (5’-3’) | Fluorescent dye (5’) | Modification (3’) |
| --- | --- | --- | --- |
| 16S Forward | CCTGGGAAACTGGGTCTAAT |  |  |
| 16S Probe | ACCGGATAGGACCACGGGATGC | FAM | ZEN (internal), IBFQ |
| 16S Reverse | CGCTTTCCACCACAAGAC |  |  |

#### **Figure S1 Half-life of 16S rRNA of 9 days is required to explain the gap between CFU and 16S rRNA counts in Mtb-infected mice treated with HRZE.**

We combined data from various experiments in which Mtb-infected Balb/c mice were treated with HRZE and the number of viable bacteria (CFU) and 16S rRNA were counted in the lungs. a. We show the change in CFU and 16S rRNA counts in lungs of Mtb-infected cells. Markers show the data from individual mice and lines are predictions of the LD model (eqns (2)-(4)). Mean CFU from lungs of mice declined in 56 days by 5.1 logs10, while 16S counts declined by only 1.5 logs10, resulting in a gap of 3.6 logs10. Other model parameters are $B\left( 0 \right)=2.50\times{10}^{7}$, $S\left( 0 \right)=4.39\times{10}^{8}$, $D(0)=0$ (assumed), $n_{0}=B(0)/S(0)=52.8$ (required by assuming D(0) = 0), $\delta_{B}=0.20/\text{day}$ (half-life 3.4 days), $\delta_{S}$= 0.076/day (half-life 9.2 days). b. Percent of the gap explained by the LD model at different times and for different assumed half-life of 16S rRNA. The values shown in the heatmap are calculated according to eqn. (6). Values in excess of 100% are indicated as 100%.

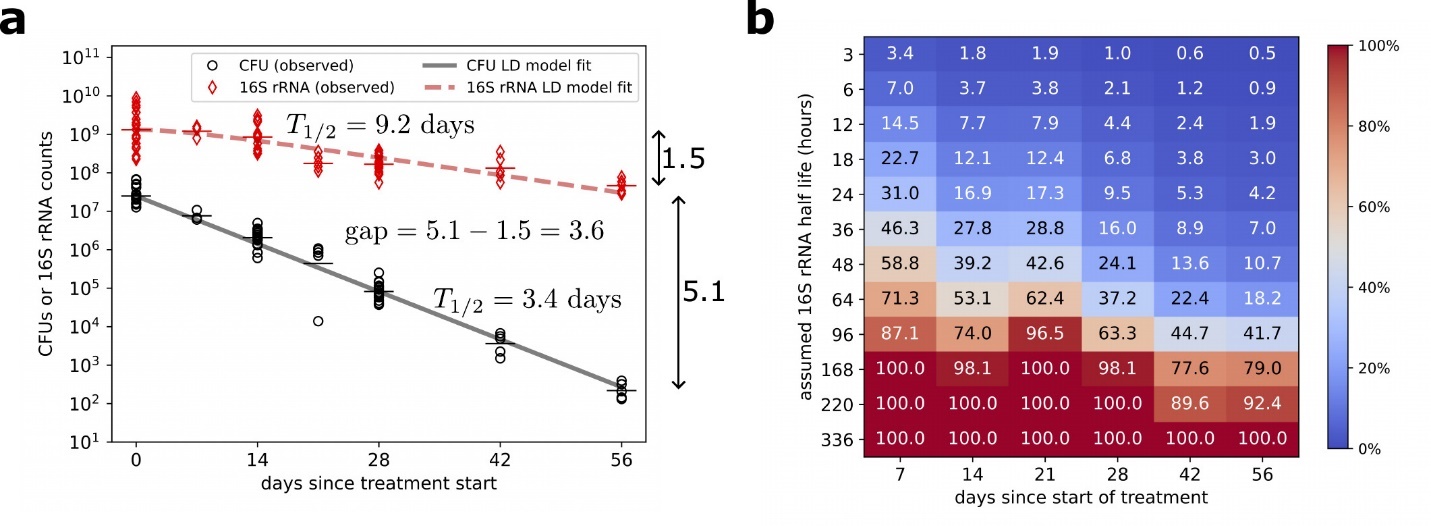

8.	Dide-Agossou C, Bauman AA, Ramey ME, et al. Combination of Mycobacterium tuberculosis RS Ratio and CFU Improves the Ability of Murine Efficacy Experiments to Distinguish between Drug Treatments. *Antimicrob Agents Chemother*. 66(4):e02310-21. doi:10.1128/aac.02310-21
